# Resting-State Electroencephalography for Continuous, Passive Prediction of Coma Recovery After Acute Brain Injury

**DOI:** 10.1101/2022.09.30.510334

**Authors:** Morteza Zabihi, Daniel B. Rubin, Sophie E. Ack, Emily J. Gilmore, Valdery Moura Junior, Sahar F. Zafar, Quanzheng Li, Michael J. Young, Brian L. Edlow, Yelena G. Bodien, Eric S. Rosenthal

## Abstract

Accurately predicting emergence from disorders of consciousness (DoC) after acute brain injury can profoundly influence mortality, acute management, and rehabilitation planning. While recent advances in functional neuroimaging and stimulus-based EEG offer the potential to enrich shared decision-making, their procedural sophistication and expense limit widespread availability or repeated performance. We investigated continuous EEG (cEEG) within a passive, “resting-state” framework to provide continuously updated predictions of DoC recovery at 24-, 48-, and 72-hour prediction horizons. To develop robust, continuous prediction models from a large population of patients with acute brain injury (ABI), we leveraged a recently described pragmatic approach transforming Glasgow Coma Scale assessment sub-score combinations into frequently assessed DoC diagnoses: coma, vegetative state, minimally conscious state with or without language, and post-injury confusional or recovered states. We retrospectively identified consecutive patients undergoing cEEG following acute traumatic brain injury (TBI), subarachnoid hemorrhage (SAH), or intracerebral hemorrhage (ICH). Models continuously predicting DoC diagnosis for multiple prediction horizons were evaluated utilizing recent clinical assessments with or without cEEG information, which comprised a comprehensive EEG feature set of 288 time, frequency, and time-frequency characteristics computed from consecutive 5-minute EEG epochs, with 6 additional features capturing each EEG feature’s temporal dynamics. Features were fed into a predictive model developed with cross-validation; the ordinal DoC diagnosis was discriminated using an ensemble of XGBoost binary classifiers. For 201 ABI patients (46 TBI, 140 SAH, 15 ICH patients comprising 27,280 cEEG-hours with concomitant clinical assessments), cEEG-augmented models accurately predicted the future DoC diagnosis at 24 hours (one-vs-rest AU-ROC, 92.4%; weighted-F1 84.1%), 48 hours (one-vs-rest AU-ROC=88%, weighted-F1=80%) and 72 hours (one-vs-rest AU-ROC=86.3%, weighted-F1=76.6%). Models were robust to utilizing different ordinal cut-points for the DoC prediction target and evaluating additional models derived from specific sub-populations using a confound-isolating cross-validation framework. The most robust features across evaluation configurations included Petrosian fractal dimension, relative power of high to low (gamma-beta to delta-alpha) EEG frequency spectra, energy within the 12-35 Hz frequency band in the short-time Fourier transform domain, and wavelet entropy. The cEEG-augmented model exceeded the performance of models using preceding clinical assessments, continuously predicting future DoC diagnosis with one-vs-rest AU-ROC in the range of 84.3-92.4% while utilizing approaches to limit overfitting. The proposed continuous, resting-state cEEG prediction method represents a promising tool to predict DoC emergence in ABI patients. Enabling these methods prospectively would represent a new paradigm of continuous prognostic monitoring for predicting coma recovery and assessing treatment response.

## Introduction

Caring for patients with coma and other disorders of consciousness (DoC) following acute brain injury (ABI) involves high-stakes decision-making compounded by diagnostic and prognostic uncertainty. Relying on clinical behavior alone to measure consciousness can result in misdiagnosis,^1,2^ leading to premature withdrawal of life-sustaining therapy or limitation of rehabilitative treatments.^3,4^ While functional MRI,^5–8^ stimulus-based EEG,^8–11^ and transcranial magnetic stimulation-EEG (TMS-EEG)^12,13^ have been demonstrated to detect consciousness and predict recovery, these tools require highly structured stimulus-based paradigms, and thus are not widely available, easily performed at scale, or repeated at a frequency required for continuous prediction.

Tools for continuously predicting coma recovery could empower clinicians and family members with updated prognostic information during the dynamic period of early recovery and secondary complications. Continuous EEG (cEEG) is a tool that offers advantages by virtue of its high temporal resolution and underlying representation of functional brain networks.^14^ As a result, cEEG has been widely utilized after ABI for cross-sectional diagnosis of DoC,^15,16^ encephalopathy,^17,18^ language dysfunction,^19,20^ and monitoring treatment response^21,22^ or neurologic deterioration.^22,23^ High-density cEEG has been employed for many of these applications,^24–28^ but has practical limitations associated with numerous collocating channels in the setting of invasive neuromonitoring devices, drains, or cranial wounds, which prevent its use in the intensive care unit (ICU) setting. Where low-density scalp coverage has been examined,^29^ no significant association was found between low-density EEG network measures and good outcomes at 3-6 months after injury, although EEG measures of functional connectivity have demonstrated promise in predicting poor outcomes in small cohorts of patients with postanoxic coma.^30^

We sought to develop models utilizing resting-state cEEG for predicting emergence from DoC continuously. Individual EEG features, such as alpha power and variability,^31,15^ have demonstrated promise for the cross-sectional classification of unresponsive wakefulness syndrome (UWS) versus minimally conscious state (MCS). We specifically aimed to develop cEEG prediction models robust to ABI etiology, age, and the time following admission, while advancing on prior investigations by incorporating ordinal rather than binary predictions, enabling continuous prediction, and including model evaluation metrics beyond the area under a receiver operating characteristic (AU-ROC) curve.^15,30^ The latter is essential in highly imbalanced classes because the AU-ROC curve may still indicate relatively high performance while misclassifying most samples from a minority class. To accomplish this, we evaluated the incremental benefit of cEEG over information from prior neurologic assessments among 201 ABI patients undergoing cEEG over a 3-year period. We developed and evaluated a multiclass, ordinal prediction model with a rolling window to facilitate predicting consciousness levels at multiple time horizons, assessing robustness using a confound-isolating cross-validation approach.

## Methods

### Study design

We performed a retrospective, single-center study of patients with ABI of different etiologies undergoing cEEG during clinical bedside monitoring of neurologic status in the Neuroscience ICU. We specified that models be constructed and tested for predicting future consciousness levels at multiple future time horizons and categorized as an ordinal outcome (i.e., coma, VS/UWS, minimally conscious state with or without language (MCS+ and MCS-), post-injury confusional state (PICS), and recovered from PICS (rPICS)). We specified sensitivity analyses examining different ordinal cut points in the level of consciousness.

### Study participants, environment, and clinical measures

We included patients aged 18 or greater, admitted to the Massachusetts General Hospital Neurosciences Intensive Care Unit between April 2016 to October 2018 with a diagnosis of traumatic brain injury (TBI), aneurysmal or non-traumatic subarachnoid hemorrhage (SAH), or intracerebral hemorrhage (ICH) who had cEEG monitoring^32^. We restricted the analysis to patients for whom EEG was initiated within 14 days of admission, recorded for at least 35 minutes, and for whom both the cEEG data and nurse-documented clinical examinations were available from our institution’s electronic data warehouse. We augmented this cohort with additional patients previously enrolled in a cohort study (May 2013-April 2016) in which these nurse-documented examinations were not yet populated in the electronic data warehouse but were instead extracted by manual chart review. Patients with other types of brain injuries, such as anoxic-ischemic injury from cardiac arrest, were excluded unless having TBI, comorbid SAH, or ICH. The institutional review board approved this retrospective study and determined that it was exempt from obtaining participants’ informed consent. Per clinical standard, cEEG was performed using the 10-20 International Standard (21 electrodes, 256 or 512 Hz sampling rate).

According to our institutional guidelines for patients with these conditions, the Glasgow Coma Scale (GCS) score^33^ was measured for patients with these conditions approximately every 2 hours; more frequent assessments were indicated for patients during periods of heightened risk, or less frequently if a patient was nearing transfer out of the intensive care environment. According to our institutional critical care nursing standards of practice, the clinical standard of care was to examine patients during sedation interruption, waiting between 5 and 20 minutes based on the nurse’s judgment.

### Prediction framework

Given an EEG time series with a lookback history of length *T*, we specified the objective to continuously predict the consciousness level of an ABI patient at a prediction horizon of *H* with a 5-minute stride. This is a rolling prediction, where each epoch and corresponding consciousness level (i.e., prediction target) were shifted forward by 5 minutes through time. We considered *T* ≤ 14 *hours* to provide the flexibility to utilize varying-length EEG recordings for up to 14 hours based on their availability at a time. For *H*, three prediction horizons of 24, 48, and 72 hours were determined to assess the effects of the prediction horizon on the performance (see Figure 1).

**Figure 1.**
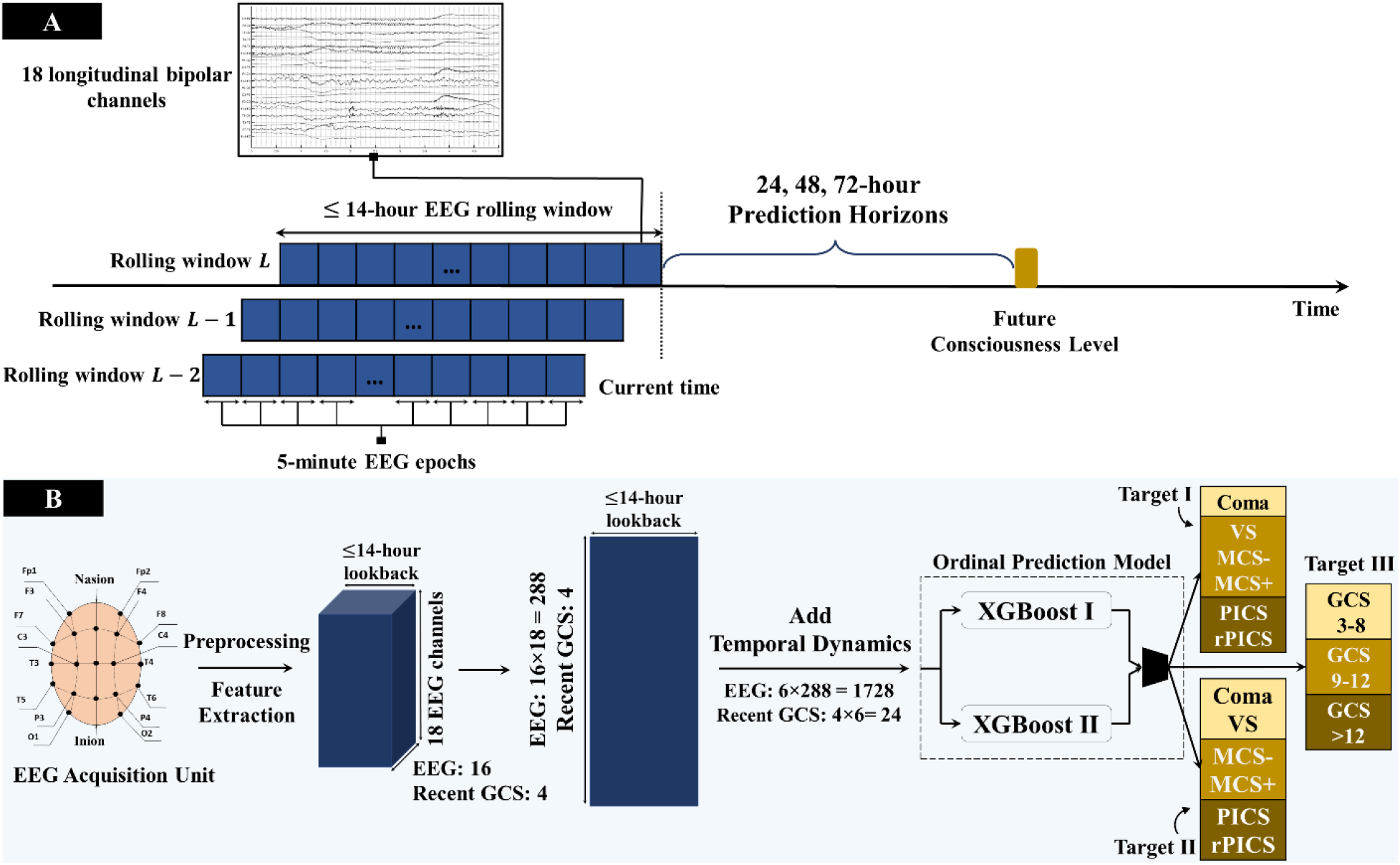
Schematic pipeline of the developed prediction approach using resting-state EEG recordings in acute brain injury patients. Future consciousness is continuously predicted using historical GCS sub-scores as a base model and historical GCS sub-scores in addition to EEG features in the augmented model, utilizing up to 14 hours of lookback data, depending on availability. **(A)** The general framework of the rolling-prediction approach with three prediction horizons of 24, 48, and 72 hours. *L* indicates the number of the rolling window with the stride of 5 minutes. **(B)** The block diagram of the proposed approach consists of preprocessing, feature extraction, and a prediction model. Six features are computed along the lookback dimension to capture the temporal dynamics of EEG features. The extracted features are fed into an ordinal prediction model formed with two XGBoost binary classifiers. The generated predictions labels *poor, moderate*, and *good* are then mapped into three ordinal target scales with clinically relevant cut-points defined based on disorders of consciousness (Targets Ⅰ & Ⅱ) and total GCS score (Target Ⅲ). MCS, minimally conscious state; PICS, post-injury confusional state; rPICS, recovered from post-injury confusional state.

### EEG feature engineering

For each patient, we utilized a longitudinal bipolar montage of scalp EEG channels containing 18 bipolar channels consisting of the following pairs: Fp1-F7, F7-T3, T3-T5, T5-O1, Fp1-F3, F3-C3, C3-P3, P3-O1, Fp2-F4, F4-C4, C4-P4, P4-O2, Fp2-F8, F8-T4, T4-T6, T6-O2, Fz-Cz, and Cz-Pz.

For preprocessing, a Butterworth bandpass filter with lower and higher cutoff frequencies of 0.5-45 Hz was applied, as well as a notch filter to remove both 60-Hz electrical noise and its 120-Hz harmonic. Data were segmented into epochs of 5 minutes with no overlap. Reported analyses were performed on these 5-minute epochs unless otherwise stated. To automatically identify possible artifactual epochs, we applied a simple rule-based method using the absolute value of EEG voltage; if any absolute value in the epoch exceeded 250 µV, the epoch was considered an artifact.^34^ EEG-based features consisted of 16 features from each of 18 bipolar channels, extracted from each 5-minute epoch. Candidate EEG features included the following:

#### Petrosian fractal dimension

There are numerous methods in a pure math setting to gauge fractal dimensions, e.g., Minkowski and Hausdorf.^35^ However, such spatial analyses are not appropriate for time series with self-affinity properties due to either the lack of well-defined special characteristics in one-dimensional signals or their relatively high computational complexity. We, therefore, utilized Petrosian’s algorithm^36,37^ to compute the time series fractal dimension as follows

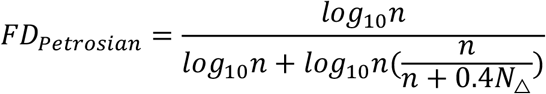

where n and N_Δ_are the length of the epoch and the number of sign changes in the first derivative of the EEG, respectively. The Petrosian fractal dimension can be considered a measure of complexity where the more complex EEG epoch leads to a higher value of its *FD*_*Petrosian*_.

#### Power spectral analysis

In many EEG classification or prediction studies, power spectral analysis in specific frequency bands is used alone or combined with other features in various tasks.^38,39^ We extracted three features from this domain. For power spectral density estimation, Welch’s method was utilized.^40^ The first feature is the relative power of 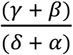, inspired by da Silveira et al.,^41^ where *β, γ, δ*, and *α* indicate the frequency bands between 12–35, >35, 1–4, and 8–12 Hz, respectively. The second and third features in this domain are the decay of the EEG power spectrum.^38,39^ We performed regression *log*_l0_*PSD* = *αlog*_10_*ω* between 4-8 and 8-12 Hz frequency bands to characterize the decay of the EEG power spectrum, *α*. Here, *PSD* and *ω* indicate the power spectral density and frequency bin in Hz, respectively.

#### Short-Time Fourier transform analysis

Short-Time Fourier Transform (STFT) characterizes time-frequency power and phase information of nonstationary signal changes over time. We computed 10 features from this domain. For STFT calculation, Welch’s method was used with a 9-second Hann window and a 6-second overlap. Also, we computed the logarithm of the amplitude spectrum as it can provide a more detailed structure while preserving the relative relation in the spectrum.^42^

The first two features are the total energy between the 4-8Hz and 12-35Hz frequency bands in the STFT domain as below

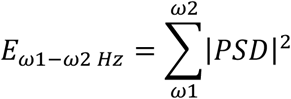

Furthermore, we extracted 8 features by tracking the prominent frequency components and their time intervals within each epoch. This is inspired by music recognition algorithms, which distill music samples into fingerprints for matching and searching.^43^ For this purpose, first, the prominent peaks were identified in each STFT time segment. These peaks indicate the highest frequency powers in the corresponding time interval. To avoid identifying too many peaks and focus on the most prominent ones, we applied a constraint to our search by specifying a minimum distance (100 bins) between the peaks. Once the peaks in all the time segments were obtained, the four most significant peaks were chosen. In addition to the four most frequent dominant frequencies, the medians of their time intervals are used as features. We computed the median of their time intervals when the same frequency component is the dominant peak in several time segments. This dynamic approach captures the most informative frequency components and their time relations from a broad spectrum rather than focusing on specific frequency bands.

#### Wavelet analysis

We extracted two features using discrete wavelet transform. We used Daubechies-4 wavelets with a decomposition level of 7. These features are based on the approximation coefficients in level 7 and detail coefficients from levels 1 to 7. The first feature is Wavelet entropy, computed as follows.^44–46^

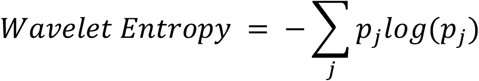

where 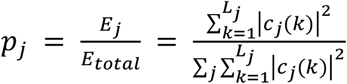, and *c*_*j*_ (*k*) is decomposition coefficients at scale *j*.

The correlation between sleep spindles and recovery of consciousness has been investigated in several studies, which is the underlying motivation for extracting the second feature in this domain.^47–50^ The frequency range associated with fast and slow spindles is 12-15 Hz.^51^ We therefore filtered the EEG signal with a passband frequency range of approximately 8-16 Hz (which may slightly change based on the sampling frequency) using wavelet decomposition. Briefly, filtering using discrete wavelet transform first decomposes the signal into approximation and detail coefficients, zeros out the details coefficients at some chosen scales, and finally assembles them back into the original signal without affecting the general shape of the signal. Once the desired passband filtering was conducted, the frequency component with maximum power spectral density was used as a feature.

### Multiclass, ordinal DoC level prediction targets

To develop continuous prediction models from a large population, we required frequent clinical assessments over a prolonged duration. We, therefore, leveraged a recently described pragmatic approach^52^ transforming repeated clinical assessments of GCS sub-scores combinations into frequently assessed DoC diagnoses: coma, VS/UWS, MCS+, MCS-, PICS, and rPICS.

Overall, we examined three targets with specific cut-points utilizing these measures. The primary ordinal prediction target (Target I) classified DoC as *poor* (coma), *moderate* (VS, MCS-, MCS+), and *good* (PICS, rPICS). In two sensitivity analyses, we examined different ordinal cut-points; Target II utilized cut-points as *poor* (Coma, VS/UWS), *moderate* (MCS-, MCS+), and *good* (PICS, rPICS), and Target III utilized cut-points based on the raw GCS scores: *poor* (total GCS= 3-8), *moderate* (total GCS=9-12), and *good* (total GCS=13-15).

### Continuous, ordinal prediction model

Our classification approach leverages discriminative models for continuous prediction of consciousness states, i.e., DoC diagnoses stratified by the criteria above, by (1) facilitating the temporal dependencies in varying length sequences, i.e., utilizing varying length historical data based on the availability of collected cEEG up to 14 hours, and (2) exploiting the ordering of information in the prediction targets rather than treating them as nominal classes.

Each 5-minute raw EEG data was transformed into a 16×18 feature space, where 16 represents the number of EEG characteristics discussed in the previous section, and 18 indicates the number of bipolar channels. Such feature space does not convey temporal information and time dependencies of adjacent epochs. To add the temporal dimension, 6 statistical features, i.e., maximum, minimum, mean, variance, 95% percentile, and interquartile range, were computed over the lookback period (up to the 14 preceding hours). This transformed each 5-minute EEG epoch into a (16×18×6) 1728-dimensional feature space with the temporal information to predict its future consciousness level, capturing each feature’s temporal dynamics and permitting discriminative methods, such as XGBoost, for sequence prediction problems. The same approach was applied to the consecutive total GCS score and its three subscale scores, i.e., 4×6 (Figure 1A).

The prediction targets, i.e., *poor, moderate*, and *good* consciousness levels, have a natural ordering which should be used to conduct a more robust analysis. However, standard classification algorithms do not often utilize ordering information in ordinal prediction and classification problems, treating the range of classes as a set of unordered values. We used an ensemble of XGBoost^53^ classifiers to account for the inherent order between classes using a previously proposed method.^54^ For the three-class classification problem, two binary XGBoost classifiers were trained. The first binary classifier was trained to estimate the probability of belonging to *moderate* and *good* classes, *Pr*(*target > poor*). The second classifier was trained to estimate the probability of belonging to the *good* class, *Pr*(*target > moderate*). Finally, the following ensemble rule was employed to generate the final prediction labels:

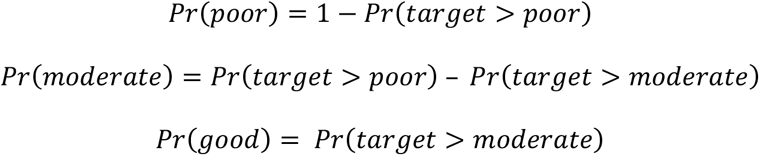

The above approach considers the ordering information and breaks down the three-class classification problem into two binary ones. We used an ensemble of 250 decision trees for each XGBoost model. No hyperparameter tuning was conducted for training to avoid the chance of overfitting, and the default values were set with learning rate=0.3, gamma=0, maximum depth=6, and the minimum child weight=1. A random under-sampling method was performed in the training phase to mitigate class imbalance.

### Model evaluation metrics

Models were evaluated for the following six metrics: accuracy, weighted- and macro-F1 scores, one-vs-one and one-vs-rest AU-ROC, and Cohen kappa measures.^55^ Furthermore, we reported confusion matrices detailing the magnitude of difference when a prediction deviated from actual. As patient-wise k-fold cross-validation was used for model assessment to avoid overfitting, we report the average of the achieved performance on the *k* held-out unseen sets.

### Selection of less-confounded features

We utilized the XGBoost feature importance values, computed as importance by information gain, for feature selection.^53^ While various techniques can quantify the relative effect of each feature on the prediction, such as Shapley additive explanations^56^ or filter methods,^57^ we chose this method as it performs the feature ranking and training in parallel, offering a relatively lower computational complexity for ranking 1728 features.

Although the main objective of this study was the “prediction” of consciousness level (and not causal interpretation), we took specific caution to diminish the effect of spurious associations on the interpretation of feature importance. As such, we utilized a previously introduced data partitioning approach^58^ to analyze the potential confounding variables in our feature relevance explanation. This “confound-isolating cross-validation” approach evaluates the feature importance where the confounding effect is absent rather than regressing out the confounding variables separately from each feature as is typical of classical statistical methods. The core idea is to choose jointly selected features across mutually exclusive partitions of data, where each partition is unique from the other by containing a dissimilar confounder distribution.

One example of evaluating robustness with confound-isolating cross-validation is for the potential confounder of etiology, i.e., TBI, SAH, or ICH. Here, the top invariant features would be calculated distinctly across each of these three etiologies. This approach ensures that feature importance is not derived by confounders that separately explain the outcome. To handle multiple confounders, we partitioned the dataset into naturally occurring strata, where each stratum is an observed combination of the confounders. This approach was taken to avoid bias due to the association between confounders and the prediction features. To address the natural occurrence that some strata may contain only a few samples, leading to underfitting the feature ranking model, we utilized patient-wise cross-validation. Specifically, the dataset was partitioned into strata, and we used all the strata except one to obtain the feature importance. We continued this process until all the strata were left out once (Figure 2A). This guaranteed enough data for the training feature ranking model and isolated one confounding stratum at each fold, albeit at the cost of *k* times training, where *k* indicates the number of strata. First, the 20 features with the highest importance from each of the two XGBoost binary classifiers were picked to find the relevant features. The built-in feature importance of the XGBoost was used for computing feature importance. Then, the common features among them were chosen as the candidate invariant features. The candidate features were fed into the feature selection approach discussed above (Figure 2) to identify the mutual features across all the confounder strata. This procedure was performed for all the determined prediction targets and horizons. While this approach evaluates models without confounders to ensure robustness in populations with any proportion of the confounder (e.g., diagnosis), risks of the approach include the potential elimination of signal and the potential for low sample size in individual partitions utilized for these deconfounding assessments.

**Figure 2.**
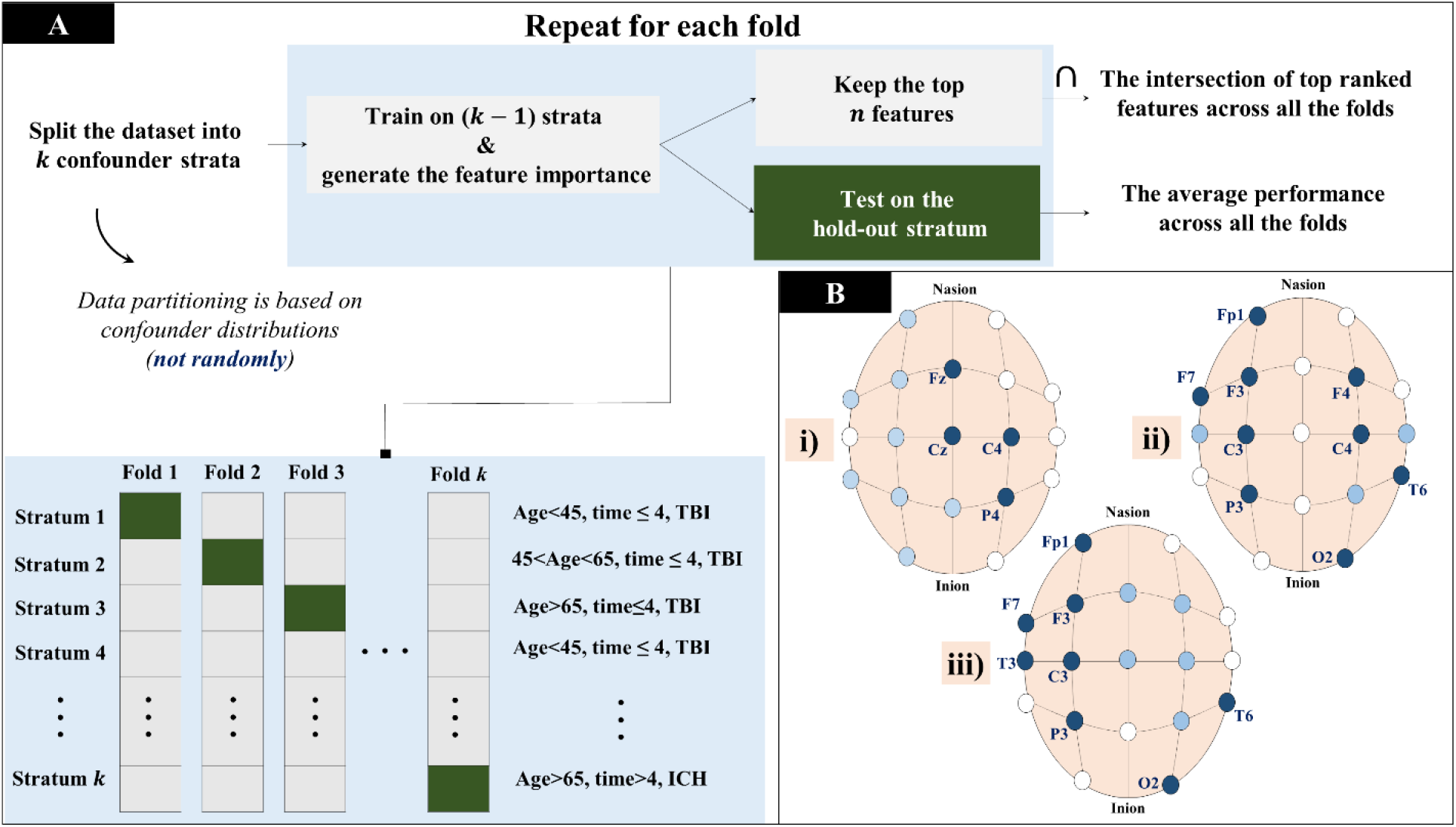
Evaluation for robustness and confounding. **(A)** Overview of the utilized framework for confound-isolating cross-validation. The three studied confounding variables are age (years old), diagnosis, and time to admission (days). *Time* refers to the time from hospital admission to EEG recording. **(B)** EEG channels from which the selected features were computed are represented for each target. The channels that were not among the selected ones are shown in white color. Channels utilized once and more than once are indicated with blue and dark blue colors, respectively. Subfigures (**i)**, (**ii)**, and (**iii)** illustrate the selected channels for prediction Targets Ⅰ, Ⅱ, and Ⅲ, respectively. TBI, traumatic brain injury

Our study focused on assessing the population by deconfounding three potential confounder variables based on the data distribution: etiology (TBI, SAH, and ICH), age (≤45, 46-65, and >65 years old), and the current monitoring time of the rolling window (≤4, >4 days following admission), 3×3×2=18 strata. Other unseen confounders, such as sedation, could affect the feature analysis. However, the purpose here was not to assess causation by exhaustively eliminating strata, but rather to promote robustness by identifying less-confounded features at the cost of each confounder stratum, further limiting the cohort size within each fold.

## Data availability

The data used in this study are available upon reasonable request, including institutional approval.

## Software

All analyses were conducted using the Python programming language. The developed feature extraction and classification are built on top of the open-source software libraries PyWavelets,^59^ SciPy,^60^ NumPy,^61^ and XGBoost.^53^

## Results

### Patient characteristics

27,280 hours of EEG recordings from 201 distinct patients with TBI, SAH, and ICH (n=46, n=140, n=15, respectively) met inclusion criteria. EEG data from nine patients with DoC were unusable either due to technical difficulties or insufficient data for the prespecified prediction horizons. Detailed patient characteristics, including the duration of recordings, are shown in Table 1. Of note, the duration of cEEG for SAH patients was longer than for TBI and ICH patients due to an institutional guideline for cEEG monitoring through the window of vasospasm or delayed cerebral ischemia risk, although the local institutional guidelines also recommend at least 24 hours of EEG for patients with TBI and ICH.

**Table 1.** Patient Characteristics.

| Patient Characteristic | TBI<br>(n=46) | SAH<br>(n=140) | ICH<br>(n=15) |
| --- | --- | --- | --- |
| Age (years), mean±SD | 61.5 ± 19.44 | 58.47 ± 13.48 | 64.93 ± 11.67 |
| Age (years), median [IQR] | 59.5 [46-80.5] | 59 [50-67] | 64 [59.5-72] |
| Weight (kg), mean±SD | 74.76 ± 0.45 | 74.99 ± 0.41 | 70.93 ± 14.24 |
| Weight (kg), median [IQR] | 70 [64.25-83.75] | 72 [68-76] | 70 [65.5-78] |
| Sex (female), n (%) | 13 (28.26%) | 108 (77.15%) | 6 (40%) |
| Latency to EEG initiation (days from admission), mean±SD | 0.86 ± 1.64 | 0.82 ± 1.55 | 1.53 ± 1.54 |
| EEG duration (hours), mean±SD | 43.67 ± 42.87 | 149.51 ± 70.14 | 61.88 ± 40.92 |
| EEG duration (hours), median [IQR] | 29.7 [20.95-43.06] | 154.25 [91.75-194.85] | 51.41 [29.91-88.7] |
| Initial GCS score, mean±SD | 8.43 ± 4.38 | 9.96 ± 3.81 | 7.8 ± 4.41 |
| Initial GCS score, median [IQR] | 8 [7-14] | 10 [5-13] | 6 [4.25-12] |
| Discharge GCS score, mean±SD | 11.69 ± 4.24 | 12.39 ± 3.35 | 11.06 ± 4.44 |
| Discharge GCS score, median [IQR] | 14 [10-15] | 14 [9-15] | 14 [8.25-15] |
| Initial DoC level, n (%) |  |  |  |
| Coma | 10 (21.73%) | 9 (6.42%) | 3 (20%) |
| VS/UWS | 0 (0%) | 2 (1.42%) | 2 (13.33%) |
| MCS - | 6 (13.04%) | 30 (21.42%) | 6 (40%) |
| MCS + | 13 (28.26%) | 24 (17.14%) | 1 (6.66%) |
| PICS | 13 (28.26%) | 28 (20%) | 1 (6.66%) |
| rPICS | 4 (8.69%) | 47 (33.57%) | 2 (13.33%) |
| Discharge DoC level, n (%) |  |  |  |
| Coma | 9 (19.56%) | 8 (5.71%) | 4 (26.66%) |
| VS/UWS | 3 (6.52%) | 15 (10.71%) | 1 (6.66%) |
| MCS - | 5 (10.86%) | 15 (10.71%) | 3 (20%) |
| MCS + | 6 (13.04%) | 16 (11.42%) | 3 (20%) |
| PICS | 15 (32.60%) | 22 (15.71%) | 3 (20%) |
| rPICS | 8 (17.39%) | 64 (45.71%) | 1 (6.66%) |

*PICS, confusional state; GCS, Glasgow Coma Scale score; MCS-, minimally conscious state without language function; MCS+, minimally conscious state with language function; rPICS, recovered from PICS; TBI, traumatic brain injury; VS/UWS, vegetative state/unresponsive wakefulness syndrome*

### Model performance for ordinal prediction targets and time horizons

To examine whether the EEG markers could improve discrimination of different consciousness levels, we compared their performance with the recent GCS in a patient-wise standard 5-fold cross-validation scheme across different prediction targets and horizons.

The developed approach obtained the highest performance when EEG features were added to the base model of recent GCS scores (Table ***2***). This added value was present across all types of prediction targets and horizons. More specifically, for Target Ⅰ the obtained one-vs-rest AU-ROC curve and macro-F1, respectively, were 92.4% and 70.2% at 24h, 88% and 64.8% at 48h, and 86.3% and 61.5% at 72h. For Target Ⅱ, the obtained one-vs-rest AU-ROC curve and macro-F1 were 92.4% and 75.7%, at 24h 88% and 70.2% at 48h, and 85.3% and 63.3% at 72h. Finally, for Target Ⅲ, one-vs-rest AU-ROC curve and macro-F1 were 90.8% and 74% at 24h, 85.7% and 65% at 48h, and 84% and 61% at 72h. Detailed evaluations of cEEG, historical GCS, and their combinations at different prediction horizons are shown in Supplementary Tables S1-S3.

**Table 2.** Performance of multiclass prediction for various time horizons and targets in a 5-fold cross-validation scheme. The results show the performance on the unseen test set data.

| Evaluation metric, mean (SD) | 24-h Horizon |  | 48-hour Horizon |  | 72-hour Horizon |  |
| --- | --- | --- | --- | --- | --- | --- |
|  | Historical GCS | Historical GCS + cEEG | Historical GCS | Historical GCS + cEEG | Historical GCS | Historical GCS + cEEG |
| *Target I* |  |  |  |  |  |  |
| Weighted-F1 | 0.773 (0.023) | 0.841 (0.026) | 0.742 (0.039) | 0.800 (0.052) | 0.701 (0.014) | 0.766 (0.026) |
| Macro-F1 | 0.645 (0.025) | 0.702 (0.044) | 0.606 (0.007) | 0.648 (0.062) | 0.561 (0.050) | 0.615 (0.035) |
| OVR AU-ROC | 0.866 (0.023) | 0.924 (0.009) | 0.820 (0.022) | 0.880 (0.029) | 0.798 (0.032) | 0.863 (0.018) |
| *Target II* |  |  |  |  |  |  |
| Weighted-F1 | 0.766 (0.017) | 0.814 (0.021) | 0.713 (0.030) | 0.764 (0.033) | 0.675 (0.027) | 0.713 (0.023) |
| Macro-F1 | 0.704 (0.012) | 0.757 (0.039) | 0.652 (0.030) | 0.702 (0.035) | 0.597 (0.051) | 0.633 (0.028) |
| OVR AU-ROC | 0.871 (0.014) | 0.924 (0.008) | 0.833 (0.026) | 0.880 (0.028) | 0.799 (0.028) | 0.853 (0.020) |
| *Target III* |  |  |  |  |  |  |
| Weighted-F1 | 0.746 (0.019) | 0.787 (0.025) | 0.689 (0.016) | 0.711 (0.022) | 0.656 (0.032) | 0.687 (0.016) |
| Macro-F1 | 0.693 (0.024) | 0.740 (0.025) | 0.631 (0.018) | 0.650 (0.027) | 0.584 (0.040) | 0.610 (0.023) |
| OVR AU-ROC | 0.882 (0.013) | 0.908 (0.016) | 0.833 (0.012) | 0.857 (0.014) | 0.796 (0.034) | 0.840 (0.013) |

*AU-ROC, area under the receiver-operating-characteristic curve; OVR, one-versus-rest; SD, standard deviation*

### Evaluation of feature robustness

The framework for feature calculation is shown in Figure 2. Feature robustness was evaluated using the previously discussed confound-isolating cross-validation. The joint features across different prediction horizons, i.e., 24, 48, and 72 hours, were chosen for each prediction target using this method are shown in Supplementary Tables S4-S6. These results show that the Petrosian fractal dimension, the relative power of 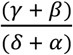, STFT energy in the 12-35 Hz frequency band, wavelet entropy, and the time interval of prominent frequency components extracted from the STFT domain were associated with the consciousness levels independent of the determined prediction targets and time horizons. As shown in Figure 2B, the selected features were extracted from the Fp1-F7, Fp1-F3, C3-P3, and T6-O2 channels more than once for prediction types Ⅱ and Ⅲ. The Petrosian fractal dimension was the most selected feature across different configurations, and none of the extracted features were selected from the Fp2-F8 channel through the confound-isolating cross-validation. Examples of features extracted from these parameters and their change in relation to the clinical trajectory are shown in Figure 3.

**Figure 3.**
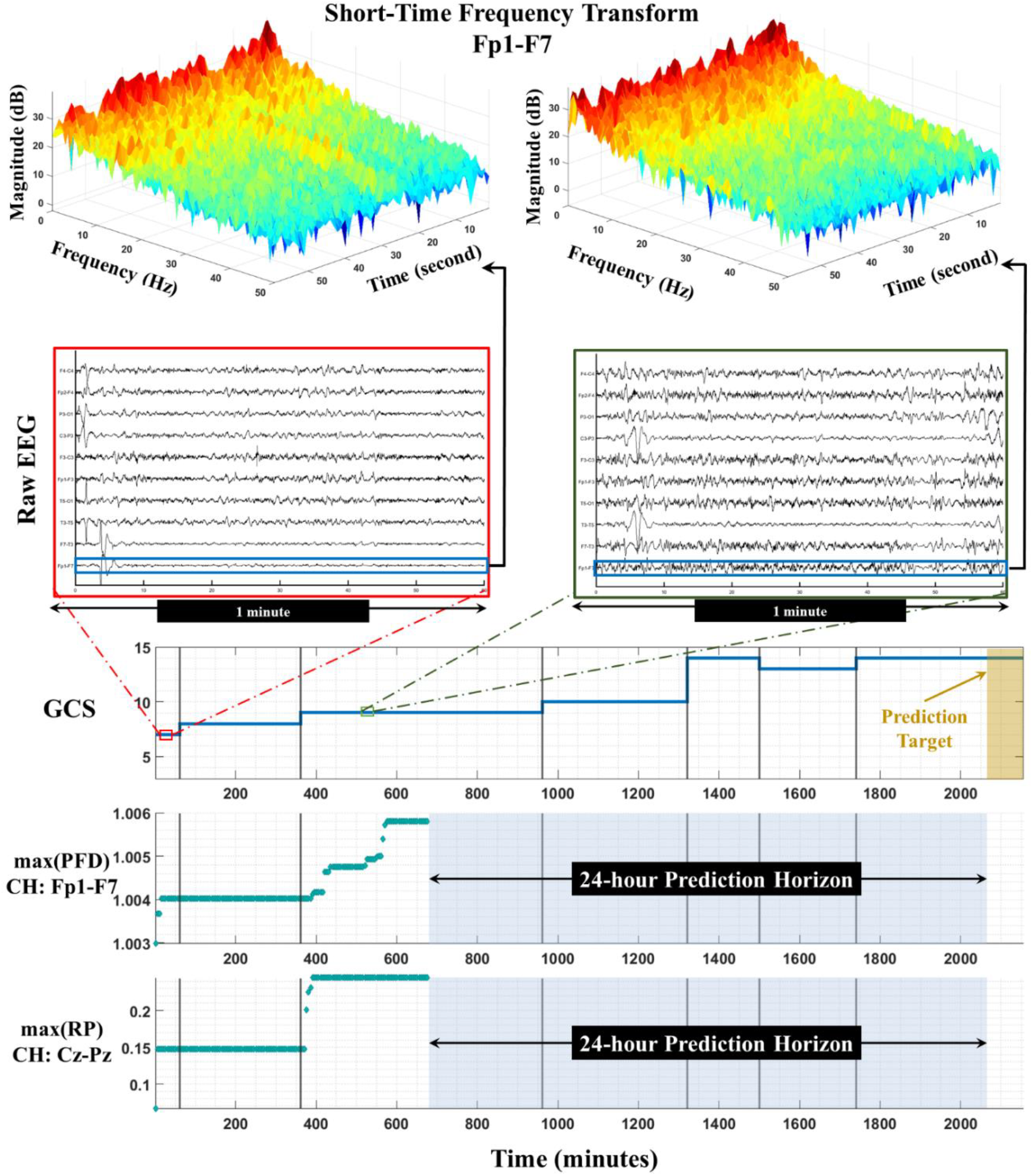
Example feature dynamics in a non-sedated TBI patient over 35 hours of cEEG. GCS, rolling maximum of the Petrosian fractal dimension (PFD) in channel Fp1-F7, rolling maximum of the relative power (RP) of 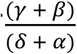 changes in channel Cz-Pz, and two random 1-minute epochs with different GCS scores with their corresponding Fp1-F7*’*s STFT are shown. As can be seen, the dynamics of the features have predictive value and are associated with the future trend of GCS. *CH, channel*.

### Evaluation of model robustness

To assess whether the selected EEG features were invariant under different confounder distributions, we evaluated the prediction model’s performance for robustness by employing the same confound-isolating approach. Figure 4 displays the three-dimensional covariate space for 24-hour prediction and the three prediction types. For TBI patients ≤45-year-old evaluated >4 days after admission, EEG markers did not improve the prediction performance across prediction Targets I, II, or III. The primary prediction target Type I (4A) was robust to other strata, whereas EEG markers did not improve the prediction performance for Targets Ⅱ and Ⅲ (Figure 4A and 4B) for ICH patients >65-year-old evaluated ≤4 days from admission.

**Figure 4.**
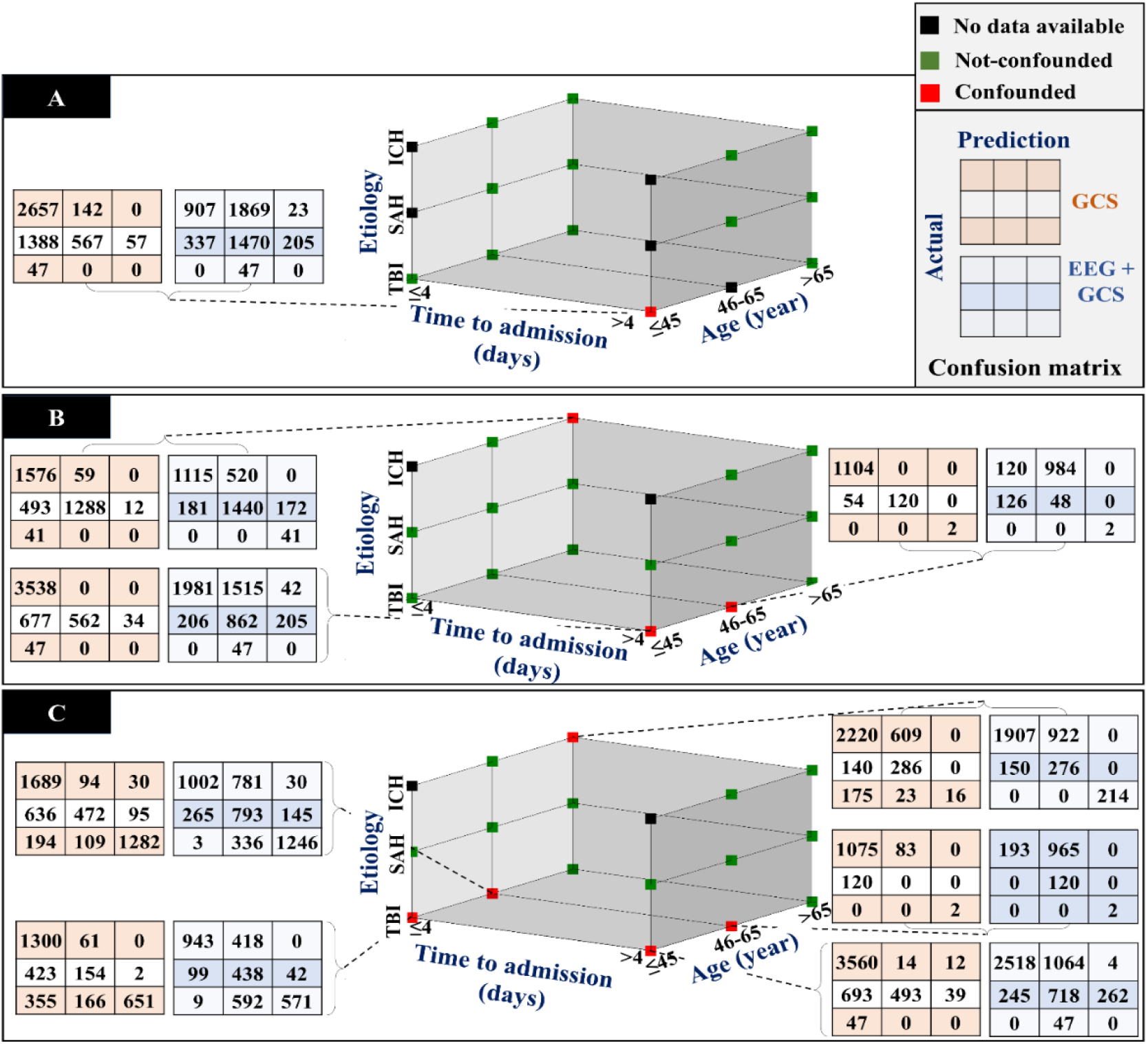
Three-dimensional confounder variables spaces for the 24-hour prediction horizon. Comparison of the obtained confusion matrices during the confound-isolating cross-validation procedure. The proposed approach and the recent GCS results are shown in blue and orange matrices, respectively. Green squares demonstrate the combinations of covariates for which cEEG improved prediction beyond that provided by historical GCS information. Red squares indicate strata in which the general prediction power of the EEG markers was less than the historical GCS. Black squares show the empty strata. Panel **(A)** corresponds to prediction Target Ⅰ, whereas panels **(B-C)** correspond to the two sensitivity analyses respectively examining prediction Targets Ⅱ and Ⅲ. Overall, most strata are green, confirming the added value of featurized cEEG information under different distributions of confounding variables. For TBI patients ≤45 years old evaluated greater than 4 days from admission, EEG markers did not augment prediction compared to using historical GCS information alone. However, the primary prediction target, Target I **(4A)** was robust to other strata. In the sensitivity analysis predicting future DoC diagnosis using different cut-points (Target II), **(Figure 4B)**, prediction performance for elderly ICH patients evaluated within 4 days of admission was also not augmented by cEEG markers. The sensitivity analysis predicting Target III (future GCS score) had the fewest strata in which cEEG markers augmented historical GCS data. In the confusion matrix, rows indicate true classes and columns indicate their corresponding prediction. The elements of the left to right (and top to bottom) show the *bad, moderate*, and *good* classes, respectively.

## Discussion

Continuous resting-state EEG features can predict coma recovery after ABI and add prognostic value compared to the recent historical trend of neurologic GCS exam scores alone. These results are robust to multiple cross-validation procedures and across various test statistics, demonstrating that EEG features convey additional predictive information beyond neurologic examination trends, even meeting this standard when developed and cross-validated across small population strata for the vast majority of potential confounder combinations.

An increasing prediction horizon only modestly diminished the performance of prediction models augmented by cEEG features, whereas prediction models utilizing historical GCS information alone decayed by a greater magnitude when extending the prediction horizon. For instance, for the primary prediction target (Target I) using only EEG, one-vs-rest AU-ROC dropped 3.9% by increasing the prediction horizon from 24h to 72h, whereas using the GCS baseline trend alone under the same conditions reduced one-vs-rest AU-ROC by 6.4%. One possible explanation is that the history of 14 hours of EEG recordings may convey more relevant information about the longer-scale dynamics of coma recovery.

Of note, *poor* recovery was the most challenging class to predict compared to *moderate* and *good* categories across all three targets with different ordinal DoC cut-points. This may be due to the cohort containing relatively few *poor* samples, approximately 7-21% of the data set (varying by target and prediction horizon). This imbalance may also explain the significant gap between weighted- and macro-F1 metrics. Additionally, predictions for Target I and Target II, future DoC ordinal rank transformed from GCS sub-score combinations, demonstrated superior performance to predictions for Target III (raw future GCS score as a total). Most GCS total scores represent a wide range of functions, and the DoC diagnoses are likely a more accurate marker of recovery.

Total GCS has an alarming rate of VS and MCS misdiagnosis compared to Coma Recovery Scale-Revised (CRS-R),^1,62,63^ and recent studies ^64^ have emphasized the imprecision of total GCS, especially in the range of 3-8 total GCS.^65^ Our predictive models included recent GCS sub-scores as features rather than total scores, which may have enhanced the ability of the GCS-only models to predict future DoC diagnosis.

One strength of these results is that the prediction approach is invariant within most confounder strata. We utilized confound-isolating cross-validation for robustness evaluation of EEG features and the proposed approach. The critical difference between confound-isolating and conventional cross-validation is randomization. In standard cross-validation, data portioning is performed randomly, but in the chosen confound-isolating approach, data is partitioned based on the confounders. While random partitioning is an excellent approach for generalization in prediction problems, randomness in cross-validation can increase the imbalance of confounding variables and should not be used for deconfounding. This confound-isolating evaluation approach demonstrated the robust association between the selected features and future consciousness level, as the selected EEG markers were invariant within small strata of different confounder combinations and also did not vary by target type or prediction horizon. Additionally, the proposed approach outperformed recent GCS even in models derived from small subsets of confounder combinations across all the prediction types.

Multiple methods intended for predicting DoC emergence have been reported, but many of these rely on technologies that are challenging to disseminate, including fMRI and high-density EEG, and most enable binary rather than ordinal prediction targets (Table 3). While it is infeasible to draw a direct comparison between the proposed coma recovery prediction methods because of different paradigms, patient characteristics, prediction horizons, evaluation methods, and metrics, the current methods overcome these limitations without sacrificing accuracy, albeit for short-term prediction targets. Should the current method be highly disseminated, patients with a high degree of residual uncertainty may benefit from additional diagnostic methods to examine concordance and provide information for long-term prognosis.

**Table 3.**
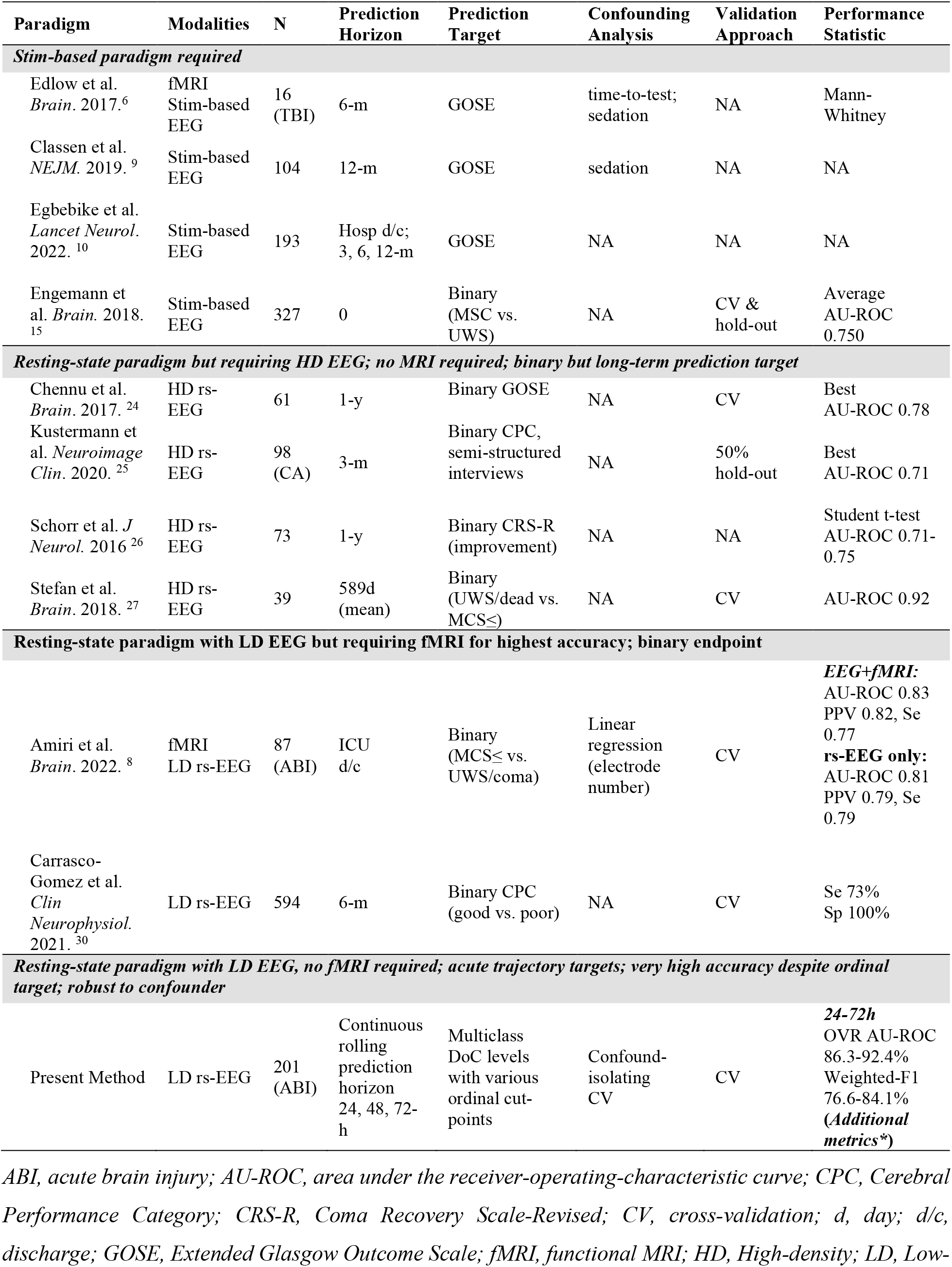
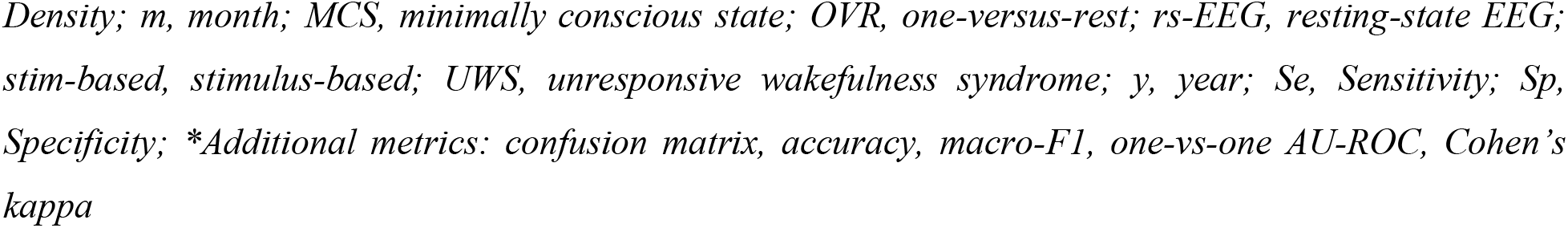
Comparison of reported methods for coma prediction.

| Paradigm | Modalities | N | Prediction Horizon | Prediction Target | Confounding Analysis | Validation Approach | Performance Statistic |
| --- | --- | --- | --- | --- | --- | --- | --- |
| ***Stim-based paradigm required*** |  |  |  |  |  |  |  |
| Edlow et al. *Brain*. 2017.<sup>6</sup> | fMRI<br>Stim-based EEG | 16 (TBI) | 6-m | GOSE | time-to-test; sedation | NA | Mann-Whitney |
| Classen et al. *NEJM*. 2019.<sup>9</sup> | Stim-based EEG | 104 | 12-m | GOSE | sedation | NA | NA |
| Egbebike et al. *Lancet Neurol*. 2022.<sup>10</sup> | Stim-based EEG | 193 | Hosp d/c; 3, 6, 12-m | GOSE | NA | NA | NA |
| Engemann et al. *Brain*. 2018.<sup>15</sup> | Stim-based EEG | 327 | 0 | Binary (MSC vs. UWS) | NA | CV & hold-out | Average AU-ROC 0.750 |
| ***Resting-state paradigm but requiring HD EEG; no MRI required; binary but long-term prediction target*** |  |  |  |  |  |  |  |
| Chennu et al. *Brain*. 2017.<sup>24</sup> | HD rs-EEG | 61 | 1-y | Binary GOSE | NA | CV | Best AU-ROC 0.78 |
| Kustermann et al. *Neuroimage Clin*. 2020.<sup>25</sup> | HD rs-EEG | 98 (CA) | 3-m | Binary CPC, semi-structured interviews | NA | 50% hold-out | Best AU-ROC 0.71 |
| Schorr et al. *J Neurol*. 2016<sup>26</sup> | HD rs-EEG | 73 | 1-y | Binary CRS-R (improvement) | NA | NA | Student t-test AU-ROC 0.71-0.75 |
| Stefan et al. *Brain*. 2018.<sup>27</sup> | HD rs-EEG | 39 | 589d (mean) | Binary (UWS/dead vs. MCS≤) | NA | CV | AU-ROC 0.92 |
| ***Resting-state paradigm with LD EEG but requiring fMRI for highest accuracy; binary endpoint*** |  |  |  |  |  |  |  |
| Amiri et al. *Brain*. 2022.<sup>8</sup> | fMRI<br>LD rs-EEG | 87 (ABI) | ICU d/c | Binary (MCS≤ vs. UWS/coma) | Linear regression (electrode number) | CV | **EEG+fMRI:**<br>AU-ROC 0.83<br>PPV 0.82, Se 0.77<br>**rs-EEG only:**<br>AU-ROC 0.81<br>PPV 0.79, Se 0.79 |
| Carrasco-Gomez et al. *Clin Neurophysiol*. 2021.<sup>30</sup> | LD rs-EEG | 594 | 6-m | Binary CPC (good vs. poor) | NA | CV | Se 73%<br>Sp 100% |
| ***Resting-state paradigm with LD EEG, no fMRI required; acute trajectory targets; very high accuracy despite ordinal target; robust to confounder*** |  |  |  |  |  |  |  |
| Present Method | LD rs-EEG | 201 (ABI) | Continuous rolling prediction horizon 24, 48, 72-h | Multiclass DoC levels with various ordinal cut-points | Confound-isolating CV | CV | **24-72h**<br>OVR AU-ROC 86.3-92.4%<br>Weighted-F1 76.6-84.1%<br>(**Additional metrics\***) |

*ABI*, acute brain injury; *AU-ROC*, area under the receiver-operating-characteristic curve; *CPC*, Cerebral Performance Category; *CRS-R*, Coma Recovery Scale-Revised; *CV*, cross-validation; *d*, day; *d/c*, discharge; *GOSE*, Extended Glasgow Outcome Scale; *fMRI*, functional MRI; *HD*, High-density; *LD*, Low-
*Density; m, month; MCS, minimally conscious state; OVR, one-versus-rest; rs-EEG, resting-state EEG; stim-based, stimulus-based; UWS, unresponsive wakefulness syndrome; y, year; Se, Sensitivity; Sp, Specificity; \*Additional metrics: confusion matrix, accuracy, macro-F1, one-vs-one AU-ROC, Cohen's kappa*

While this study is limited by its single-center design, using readily available data from existing clinical workflows allows for future external validation as it does not require stimulus-based paradigms and uses the novel approach to leverage frequently performed nurse assessments to derive DoC levels from combinations of GCS sub-scores. Additionally, while imbalance in the number of TBI, SAH, and ICH patients in the cohort represents a potential limitation, the deconfounding analysis did demonstrate robustness to etiology. We recognize that the confound-isolating cross-validation approach employed has two significant shortcomings. First, only the time-invariant confounders can be adjusted using this approach, such that dynamic ICU covariates cannot be evaluated for confounding using this method.^66^ Nevertheless, the DoC level was routinely ascertained off sedation per institutional guideline. Second, this confound-isolating approach requires categorical bins for continuous confounders such as age, such that residual confounding can be preserved within each category.^67,68^ We elected to use the clinical neurological exam rather than mortality as a prediction target to avoid self-fulfilling prophesies related to targets such as mortality, which are susceptible to withdrawal of care. Additionally, the emphasis here was on prediction, such that confounder analyses were intended to assess robustness and generalizability rather than define causal relationships.

The feature engineering included handcrafted features, i.e., nominated using human expertise, with an ensemble of XGBoosts for prediction modeling. We choose this approach as various studies have demonstrated that end-to-end deep learning approaches, with purely data-driven feature learning, perform significantly lower than expectations.^69,70^ Although the performance of any machine learning model depends highly on feature engineering, hyperparameter tuning, and the data generation process, several empirical studies have shown that these approaches to feature engineering can be highly valuable across a variety of applications when paired with the XGBoost framework.^71,72^ One future direction to examine additional benefits of deep learning would be to facilitate models combining these approaches. As previously mentioned, we did not conduct hyperparameter tunning across the XGBoost cross-validation folds to avoid overfitting.

Although nested cross-validation is an effective solution in general for hyperparameter estimation and provides a reliable generalization performance, it can increase the bias in a dataset with severe confounder imbalance. For instance, in our cohort, ICH was less common in the cohort than other diagnoses. With a severely skewed confounder distribution, if we had randomly split the data into training and test sets in the cross-validation scheme, the confounder distribution in most folds likely would have inherited the same etiology distribution from the original data. As a result, only a few samples in the training data of the most folds would potentially belong to ICH patients, because the data partitioning is performed with a uniform probability distribution.^73^ Often, machine learning approaches hypothesize that training and test sets have the same distribution, but such concerns should be cautiously approached, particularly in clinical applications. We specified our approach accordingly to promote the robustness and rigor of the methods.

Other approaches taken to ensure rigor and robustness included patient-wise evaluation and temporal ordering. Patient-wise evaluation avoids data leakage that could occur if different samples of the same patient were present in both a training and testing partition. We took particular care to obey the autoregressive property by maintaining the chronological order of EEG sequences. For predicting the consciousness level in *H*, the model only relies on cEEG data from the present period preceding the time horizon (*t − H*), looking backward through a fixed historical period (*t − T − H*). More history of EEG becomes available as we move forward through time, leading to more accurate predictions. Although our design limiting the historical lookback period has the potential for a hugely adverse effect on the evaluation performance, this rolling prediction is essential for the realistic evaluation of continuous prediction models, especially for real-time applications.

Important future directions will include validating in external data sets, examining how intercurrent changes in treatment affect predictions, and evaluating how the implementation of real-time continuous prediction influences shared decision making. For example, the approach might encourage more aggressive rehabilitation strategies, appeals for insurance coverage, and short-term trials of therapy when a high probability of coma recovery is predicted. Predicting poor outcomes may have a different effect on decision-making, either because of the irreversible nature of withdrawal of care or because model performance excels at predicting good and moderate outcomes more so than predicting poor outcomes.

Overall, these methods offer the potential for continuous prognostic monitoring (Figure 5). Conceptually, this new approach leverages continuously accruing clinical and EEG data so that updated predictions of DoC trajectory can be considered in context during iterative shared decision making. The methods inherently promote a continuous dialogue because predictions utilizing early data, despite a high accuracy in this study, can change with updated electroclinical information.

**Figure 5.**
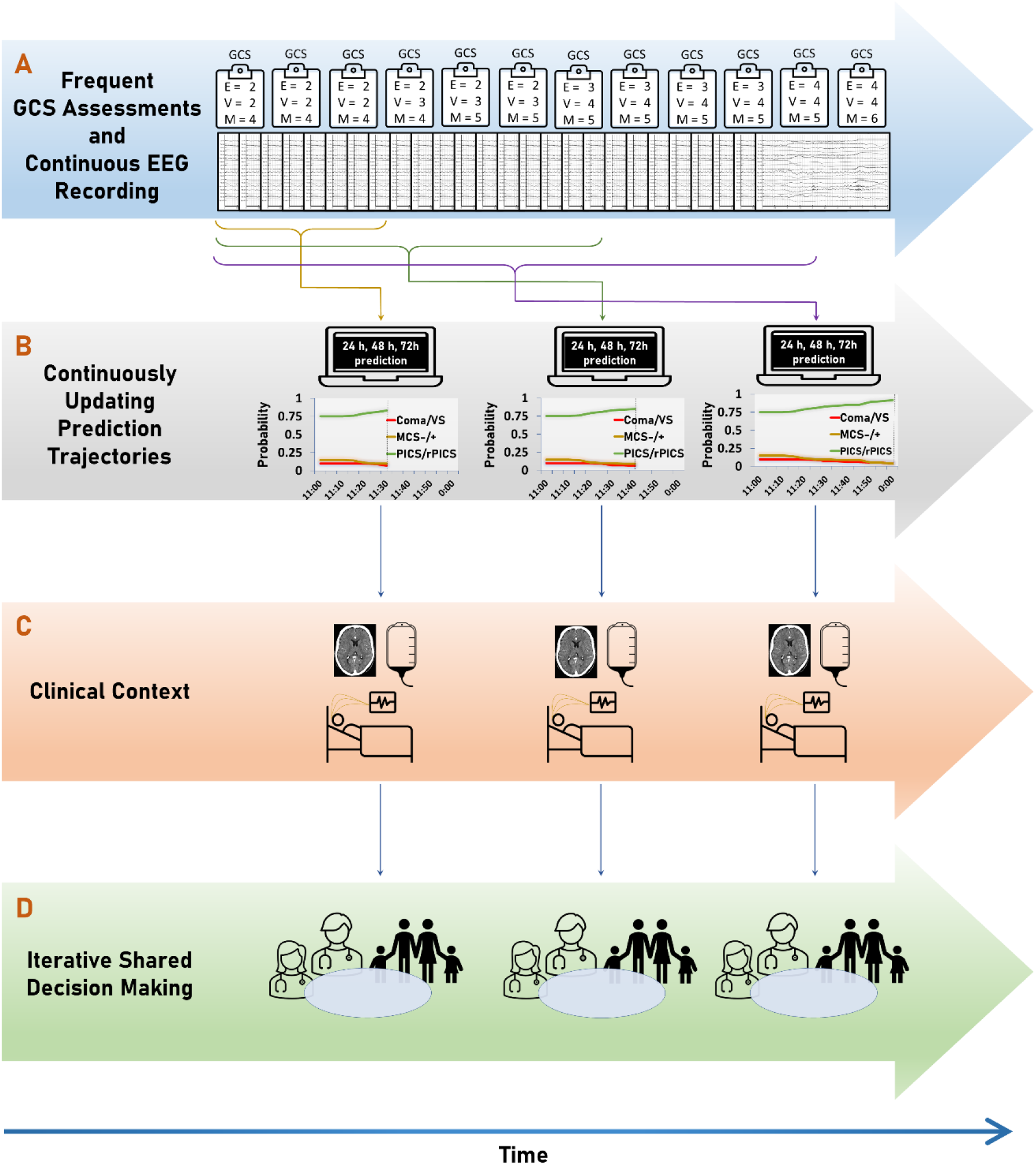
Conceptual continuous prognostic monitoring paradigm. **A)** Prognostic information from prior clinical trends on the Glasgow Coma Scale and cEEG is utilized for developing a new prediction of coma recovery every 5 minutes. **B)** Intermittently, these predictions of the patient’s recovery trajectory over the next 24, 48, and 72 hours are computed to ordinal DoC categories (e.g., Coma/VS-UWS, MCS+/MCS-, and PICS/rPICS) are available for **C)** integration within the clinical context, and **D)** shared decision making. *cEEG, continuous EEG; DoC, disorders of consciousness; MCS, minimally conscious state; PICS, post-injury confusional state (PICS); VS-UWS, vegetative state - unresponsive wakefulness syndrome*.

## Conclusion

A systematic approach using passive, resting-state low-density EEG, accurately predicted future consciousness state in ABI patients at 24-, 48-, and 72-hour time horizons, improving the performance over historical clinical assessments alone. By achieving ROC-AUC in the range of 84%-92.4% for each ordinal DoC diagnosis versus others for a range of configurations, we showed that the predictive power of the EEG-based features not only improves the prediction performance but also reveals additional information compared to clinical baseline information. These EEG features were robust to a rigorous approach to reduce confounding. While predicting a *poor* outcome remains most challenging, predictions of improvement (i.e., good outcomes) are most likely to change practice by encouraging time-limited treatment trials. While further studies should evaluate the effect of treatment strategies on the evolution of consciousness trajectory as well as cross-institutional performance, the continuous nature of the proposed approach allows for real-time application, which can potentially capture the impact of treatment in real-time.

## Abbreviations

TMS: transcranial magnetic stimulation
GCS: Glasgow Coma Scale
ABI: acute brain injury
ICU: intensive care unit
DoC: disorders of consciousness
VS/UWS: vegetative state / unresponsive wakefulness syndrome
MCS + / -: minimally conscious state with / without language
PICS: post-injury confusional state
r-PICS: recovered PICS
TBI: traumatic brain injury
SAH: subarachnoid hemorrhage
ICH: intracerebral hemorrhage
AU-ROC: the area under a receiver operating characteristic
STFT: Short-Time Fourier Transform
OVR: one-vs-rest
ML: machine learning
DL: deep learning
CRS-R: Coma Recovery Scale-Revised

## Supplementary material

Supplementary material is available as attached.

