## Supplementary materials for "Resting-State Electroencephalography for Continuous, Passive Prediction of Coma Recovery After Acute Brain Injury"

### Supplementary material

**Supplementary Table S1. Performance of the multiclass prediction for Target I.** The average (standard deviation) prediction results of the unseen data in a patient-wise 5-fold cross-validation scheme for the Target I, i.e., *poor* (Coma), *moderate* (VS, MCS-, MCS+), and *good* (PTCS, rPTCS)

| | **Evaluation metric** | **24 hr.** | | | **48 hr.** | | | **72 hr.** | | |
|---|---|---|---|---|---|---|---|---|---|---|
| **cEEG** | Accuracy | 0.765 (0.015) | | | 0.726 (0.045) | | | 0.703 (0.027) | | |
| | Weighted-F1 | 0.753 (0.019) | | | 0.713 (0.053) | | | 0.693 (0.029) | | |
| | Macro-F1 | 0.574 (0.04) | | | 0.565 (0.066) | | | 0.549 (0.035) | | |
| | One-vs-one AU-ROC | 0.777 (0.026) | | | 0.767 (0.053) | | | 0.756 (0.039) | | |
| | One-vs-rest AU-ROC | 0.832 (0.01) | | | 0.8 (0.05) | | | 0.793 (0.03) | | |
| | Cohen kappa | 0.557 (0.024) | | | 0.492 (0.075) | | | 0.447 (0.041) | | |
| | Confusion[1] matrix | 493 | 3012 | 458 | 412 | 2801 | 476 | 463 | 2284 | 582 |
| | | 758 | 19375 | 4201 | 398 | 17152 | 4696 | 471 | 14427 | 5506 |
| | | 125 | 4970 | 23892 | 117 | 6289 | 21628 | 118 | 5867 | 20793 |
| **GCS** | Accuracy | 0.737 (0.025) | | | 0.706 (0.033) | | | 0.662 (0.018) | | |
| | Weighted-F1 | 0.773(0.023) | | | 0.742 (0.039) | | | 0.701 (0.014) | | |
| | Macro-F1 | 0.645 (0.025) | | | 0.606 (0.007) | | | 0.561 (0.05) | | |
| | One-vs-one AU-ROC | 0.856 (0.026) | | | 0.811 (0.017) | | | 0.792 (0.032) | | |
| | One-vs-rest AU-ROC | 0.866 (0.023) | | | 0.820 (0.022) | | | 0.798 (0.032) | | |
| | Cohen kappa | 0.57 (0.025) | | | 0.522 (0.044) | | | 0.45 (0.046) | | |
| | Confusion matrix | 3058 | 857 | 49 | 2540 | 1101 | 48 | 2344 | 915 | 69 |
| | | 7428 | 15118 | 1788 | 6860 | 13211 | 2174 | 6992 | 10556 | 2856 |
| | | 2920 | 1977 | 24090 | 3152 | 2542 | 22340 | 3399 | 2796 | 20582 |
| **cEEG + GCS** | Accuracy | 0.848 (0.021) | | | 0.8121 (0.045) | | | 0.774 (0.024) | | |
| | Weighted-F1 | 0.841 (0.026) | | | 0.8 (0.052) | | | 0.766 (0.026) | | |
| | Macro-F1 | 0.702 (0.044) | | | 0.648 (0.062) | | | 0.615 (0.035) | | |
| | One-vs-one AU-ROC | 0.889 (0.021) | | | 0.842 (0.03) | | | 0.815 (0.039) | | |
| | One-vs-rest AU-ROC | 0.924 (0.009) | | | 0.88 (0.029) | | | 0.863 (0.018) | | |
| | Cohen kappa | 0.719 (0.035) | | | 0.651 (0.08) | | | 0.582 (0.037) | | |
| | Confusion matrix | 1136 | 2695 | 133 | 675 | 2886 | 127 | 618 | 2534 | 177 |
| | | 717 | 21694 | 1923 | 480 | 19215 | 2549 | 737 | 16157 | 3510 |
| | | 8 | 3237 | 25742 | 57 | 4069 | 23907 | 91 | 4243 | 22444 |

[1] In the confusion matrix, rows indicate true classes and columns indicate their corresponding prediction. Left to right (and top to bottom) elements show the *bad*, *moderate*, and *good* classes, respectively. The reported confusion matrices in this article are the average of the obtained confusion matrices over the unseen hold-out folds.

**Supplementary Table S2. Performance of the multiclass prediction for Target II.** The average (standard deviation) prediction results of the unseen data in a patient-wise 5-fold cross-validation scheme for the Target II, i.e., *poor* (Coma, VS), *moderate* (MCS-, MCS+), and *good* (PTCS, rPTCS).

<table>
<tr><th></th><th>Evaluation metric</th><th colspan="3">24 hr.</th><th colspan="3">48 hr.</th><th colspan="3">72 hr.</th></tr>
<tr><td rowspan="9">cEEG</td><td>Accuracy</td><td colspan="3">0.694 (0.037)</td><td colspan="3">0.654 (0.05)</td><td colspan="3">0.633 (0.018)</td></tr>
<tr><td>Weighted-F1</td><td colspan="3">0.677 (0.0502)</td><td colspan="3">0.641 (0.054)</td><td colspan="3">0.623 (0.022)</td></tr>
<tr><td>Macro-F1</td><td colspan="3">0.5771 (0.03)</td><td colspan="3">0.541 (0.054)</td><td colspan="3">0.522 (0.024)</td></tr>
<tr><td>One-vs-one AU-ROC</td><td colspan="3">0.79 (0.028)</td><td colspan="3">0.753 (0.043)</td><td colspan="3">0.742 (0.015)</td></tr>
<tr><td>One-vs-rest AU-ROC</td><td colspan="3">0.822 (0.028)</td><td colspan="3">0.786 (0.041)</td><td colspan="3">0.77 (0.013)</td></tr>
<tr><td>Cohen kappa</td><td colspan="3">0.479 (0.035)</td><td colspan="3">0.411 (0.072)</td><td colspan="3">0.368 (0.015)</td></tr>
<tr><td rowspan="3">Confusion matrix</td><td>1467</td><td>6112</td><td>828</td><td>1400</td><td>5451</td><td>1155</td><td>1464</td><td>4708</td><td>1483</td></tr>
<tr><td>1715</td><td>14566</td><td>3609</td><td>1813</td><td>12139</td><td>3975</td><td>1869</td><td>9623</td><td>4586</td></tr>
<tr><td>222</td><td>4907</td><td>23857</td><td>330</td><td>5917</td><td>21787</td><td>330</td><td>5479</td><td>20968</td></tr>
<tr><td rowspan="9">GCS</td><td>Accuracy</td><td colspan="3">0.752 (0.019)</td><td colspan="3">0.701 (0.029)</td><td colspan="3">0.663 (0.031)</td></tr>
<tr><td>Weighted-F1</td><td colspan="3">0.766 (0.017)</td><td colspan="3">0.713 (0.03)</td><td colspan="3">0.675 (0.027)</td></tr>
<tr><td>Macro-F1</td><td colspan="3">0.704 (0.012)</td><td colspan="3">0.652 (0.03)</td><td colspan="3">0.597 (0.051)</td></tr>
<tr><td>One-vs-one AU-ROC</td><td colspan="3">0.865 (0.016)</td><td colspan="3">0.826 (0.022)</td><td colspan="3">0.79 (0.028)</td></tr>
<tr><td>One-vs-rest AU-ROC</td><td colspan="3">0.871 (0.014)</td><td colspan="3">0.833 (0.026)</td><td colspan="3">0.799 (0.028)</td></tr>
<tr><td>Cohen kappa</td><td colspan="3">0.608 (0.022)</td><td colspan="3">0.527 (0.044)</td><td colspan="3">0.455 (0.061)</td></tr>
<tr><td rowspan="3">Confusion matrix</td><td>6929</td><td>1411</td><td>67</td><td>6108</td><td>1780</td><td>119</td><td>5640</td><td>1789</td><td>225</td></tr>
<tr><td>5932</td><td>12143</td><td>1816</td><td>6156</td><td>9437</td><td>2334</td><td>5815</td><td>7142</td><td>3122</td></tr>
<tr><td>2994</td><td>1952</td><td>24040</td><td>3184</td><td>2547</td><td>22303</td><td>3268</td><td>2686</td><td>20824</td></tr>
<tr><td rowspan="9">cEEG + GCS</td><td>Accuracy</td><td colspan="3">0.815 (0.02)</td><td colspan="3">0.764 (0.033)</td><td colspan="3">0.713 (0.021)</td></tr>
<tr><td>Weighted-F1</td><td colspan="3">0.814 (0.021)</td><td colspan="3">0.764 (0.033)</td><td colspan="3">0.713 (0.023)</td></tr>
<tr><td>Macro-F1</td><td colspan="3">0.757 (0.039)</td><td colspan="3">0.702 (0.035)</td><td colspan="3">0.633 (0.028)</td></tr>
<tr><td>One-vs-one AU-ROC</td><td colspan="3">0.907 (0.017)</td><td colspan="3">0.855 (0.033)</td><td colspan="3">0.822 (0.023)</td></tr>
<tr><td>One-vs-rest AU-ROC</td><td colspan="3">0.924 (0.008)</td><td colspan="3">0.88 (0.028)</td><td colspan="3">0.853 (0.02)</td></tr>
<tr><td>Cohen kappa</td><td colspan="3">0.683 (0.042)</td><td colspan="3">0.603 (0.051)</td><td colspan="3">0.51 (0.045)</td></tr>
<tr><td rowspan="3">Confusion matrix</td><td>4477</td><td>3771</td><td>159</td><td>3771</td><td>4002</td><td>233</td><td>3048</td><td>4242</td><td>365</td></tr>
<tr><td>1712</td><td>16556</td><td>1623</td><td>1815</td><td>13694</td><td>2419</td><td>2150</td><td>10544</td><td>3384</td></tr>
<tr><td>64</td><td>3183</td><td>25739</td><td>150</td><td>4109</td><td>23775</td><td>208</td><td>4009</td><td>22561</td></tr>
</table>

**Supplementary Table S3. Performance of the multiclass prediction for Target III.** The average (standard deviation) prediction results of the unseen data in a patient-wise 5-fold cross-validation scheme for the Target III, i.e., *poor* (total GCS= 3-8), *moderate* (total GCS=9-12), and *good* (total GCS=13-15)

| | **Evaluation metric** | **24 hr.** | | | **48 hr.** | | | **72 hr.** | | |
|---|---|---|---|---|---|---|---|---|---|---|
| **cEEG** | Accuracy | 0.655 (0.05) | | | 0.609 (0.049) | | | 0.6058 (0.034) | | |
| | Weighted F1 | 0.65 (0.055) | | | 0.604 (0.056) | | | 0.599 (0.029) | | |
| | Macro F1 | 0.584 (0.045) | | | 0.532 (0.052) | | | 0.513 (0.048) | | |
| | One-vs-one AU-ROC | 0.776 (0.032) | | | 0.741 (0.035) | | | 0.729 (0.035) | | |
| | One-vs-rest AU-ROC | 0.806 (0.027) | | | 0.772 (0.032) | | | 0.762 (0.028) | | |
| | Cohen kappa | 0.444 (0.056) | | | 0.363 (0.058) | | | 0.338 (0.051) | | |
| | Confusion matrix | 3771 | 7046 | 1682 | 2944 | 6424 | 1908 | 2403 | 5459 | 2198 |
| | | 3177 | 9868 | 2958 | 3004 | 8280 | 3417 | 2271 | 7385 | 4017 |
| | | 573 | 4394 | 23817 | 600 | 5692 | 21698 | 558 | 5364 | 20856 |
| **GCS** | Accuracy | 0.74 (0.018) | | | 0.684 (0.014) | | | 0.651 (0.034) | | |
| | Weighted F1 | 0.746 (0.019) | | | 0.689 (0.016) | | | 0.656 (0.032) | | |
| | Macro F1 | 0.693 (0.024) | | | 0.631 (0.018) | | | 0.584 (0.04) | | |
| | One-vs-one AU-ROC | 0.865 (0.011) | | | 0.812 (0.011) | | | 0.776 (0.03) | | |
| | One-vs-rest AU-ROC | 0.882 (0.013) | | | 0.833 (0.012) | | | 0.796 (0.034) | | |
| | Cohen kappa | 0.588 (0.028) | | | 0.5 (0.029) | | | 0.44 (0.055) | | |
| | Confusion matrix | 10159 | 2100 | 239 | 8366 | 2423 | 488 | 7190 | 2271 | 600 |
| | | 6032 | 8345 | 1626 | 6375 | 6375 | 2047 | 5929 | 4900 | 2843 |
| | | 2985 | 1864 | 23935 | 3371 | 2346 | 22274 | 3651 | 2161 | 20965 |
| **cEEG + GCS** | Accuracy | 0.783 (0.024) | | | 0.708 (0.024) | | | 0.684 (0.014) | | |
| | Weighted F1 | 0.787 (0.025) | | | 0.711 (0.022) | | | 0.687 (0.016) | | |
| | Macro F1 | 0.74 (0.025) | | | 0.65 (0.027) | | | 0.61 (0.023) | | |
| | One-vs-one AU-ROC | 0.888 (0.014) | | | 0.827 (0.016) | | | 0.807 (0.014) | | |
| | One-vs-rest AU-ROC | 0.908 (0.016) | | | 0.857 (0.014) | | | 0.84 (0.013) | | |
| | Cohen kappa | 0.651 (0.035) | | | 0.525 (0.024) | | | 0.474 (0.038) | | |
| | Confusion matrix | 7576 | 4504 | 419 | 5286 | 5555 | 436 | 4284 | 5018 | 759 |
| | | 2792 | 11626 | 1585 | 3166 | 9162 | 2372 | 2806 | 7918 | 2949 |
| | | 144 | 3033 | 25607 | 316 | 3900 | 23774 | 552 | 3766 | 22460 |

**Supplementary Table S4. Selected features for Target I.** Invariant features across all the confounder strata and prediction horizons for Target I.[2]

| Temporal Feature | EEG Feature | Channel |
|---|---|---|
| maximum | Petrosian fractal dimension | Fp1-F7 |
| maximum | Petrosian fractal dimension | T5-O1 |
| maximum | Relative power $\frac{(\gamma+\beta)}{(\delta+\alpha)}$ | Fz-Cz |
| maximum | Relative power $\frac{(\gamma+\beta)}{(\delta+\alpha)}$ | Cz-Pz |
| maximum | The median of the time interval of the $2^{nd}$ most frequent dominant frequency | C3-P3 |
| maximum | Wavelet entropy | C4-P4 |
| 95% percentile | STFT energy between 12-35 Hz | C4-P4 |
| 95% percentile | The median of the time interval of the $2^{nd}$ most frequent dominant frequency | Fp1-F3 |
| 95% percentile | The $1^{st}$ most frequent dominant frequency | Fz-Cz |

[2] The common features and EEG channels across all targets, prediction horizons, and confounder strata are highlighted in **Supplementary Tables S4-6**.

**Supplementary Table S5 Selected features for Target II.** Invariant features across all the confounder strata and prediction horizons for Target II

| Temporal Feature | EEG Feature | Channel |
|---|---|---|
| maximum | Petrosian fractal dimension | Fp1-F7 |
| maximum | Petrosian fractal dimension | F7-T3 |
| maximum | Petrosian fractal dimension | T6-O2 |
| maximum | Relative power $\frac{(\gamma+\beta)}{(\delta+\alpha)}$ | Fp1-F7 |
| maximum | STFT energy between 8-12 Hz | P4-O2 |
| maximum | The median of the time interval of the 1st most frequent dominant frequency | C3-P3 |
| maximum | The median of the time interval of the 2nd most frequent dominant frequency | F3-C3 |
| maximum | The median of the time interval of the 2nd most frequent dominant frequency | C3-P3 |
| maximum | The 1st most frequent dominant frequency | F4-C4 |
| maximum | The 1st most frequent dominant frequency | T6-O2 |
| maximum | The 2nd most frequent dominant frequency | F4-C4 |
| maximum | Wavelet entropy | C4-P4 |
| minimum | Petrosian fractal dimension | T4-T6 |
| minimum | STFT energy between 12-35 Hz | F4-C4 |
| 95% percentile | The median of the time interval of the 2nd most frequent dominant frequency | Fp1-F3 |
| 95% percentile | The median of the time interval of the 2nd most frequent dominant frequency | F3-C3 |

**Supplementary Table S6 Selected features for Target III.** Invariant features across all the confounder strata and prediction horizons for Target III.

| Temporal Feature | EEG Feature | Channel |
|---|---|---|
| maximum | Petrosian fractal dimension | Fp1-F7 |
| maximum | Petrosian fractal dimension | T6-O2 |
| maximum | Relative power $\frac{(\gamma+\beta)}{(\delta+\alpha)}$ | Fp1-F7 |
| maximum | STFT energy between 12-35 Hz | F3-C3 |
| maximum | The median of the time interval of the 2$^{nd}$ most frequent dominant frequency | F3-C3 |
| maximum | The median of the time interval of the 2$^{nd}$ most frequent dominant frequency | C3-P3 |
| maximum | The 4$^{th}$ most frequent dominant frequency | F7-T3 |
| maximum | Wavelet entropy | T6-O2 |
| minimum | Petrosian fractal dimension | F7-T3 |
| minimum | Relative power $\frac{(\gamma+\beta)}{(\delta+\alpha)}$ | Fz-Cz |
| variance | STFT energy between 12-35 Hz | F4-C4 |
| 95% percentile | Petrosian fractal dimension | F7-T3 |
| 95% percentile | The 3$^{rd}$ most frequent dominant frequency | C4-P4 |
| 95% percentile | The 4$^{th}$ most frequent dominant frequency | C3-P3 |
